# DNA polymerase β fingers movement revealed by single-molecule FRET suggests a partially closed conformation as a fidelity checkpoint

**DOI:** 10.1101/306175

**Authors:** Carel Fijen, Mariam Mahmoud, Rebecca Kaup, Jamie Towle-Weicksel, Joann Sweasy, Johannes Hohlbein

**Author notes:** Present Address: Jamie Towle-Weicksel, Physical Sciences Department, Rhode Island College, Providence, RI, 02908, USA.

## Abstract

The eukaryotic DNA polymerase β plays an important role in cellular DNA repair as it fills gaps in single nucleotide gapped DNA that result from removal of damaged bases. Since defects in DNA repair may lead to cancer and genetic instabilities, Pol β has been extensively studied, especially substrate binding and a fidelity-related conformational change called “fingers closing”. Here, we applied single-molecule Förster resonance energy transfer to study the conformational dynamics of Pol β. Using an acceptor labelled polymerase and a donor labelled DNA substrate, we measured distance changes associated with DNA binding and fingers movement. Our findings suggest that Pol β does not bend its gapped DNA substrate to the extent related crystal structures indicate: instead, bending seems to be significantly less profound. Furthermore, we visualized dynamic fingers closing in single Pol β-DNA complexes upon addition of complementary nucleotides and derived rates of conformational changes. Additionally, we provide evidence that the fingers close only partially when an incorrect nucleotide is bound. This “ajar” conformation found in Pol β, a polymerase of the X-family, suggests the existence of an additional fidelity checkpoint similar to what has been previously proposed for a member of the A-family, the bacterial DNA polymerase I.

## Introduction

DNA repair is pivotal for maintaining genome integrity (1). Among the most common damages of DNA are base lesions, in which the chemical structure of a single base has been altered (2, 3). These modifications may disturb proper base pairing and can lead to harmful mutations in the genome. In eukaryotes, the base excision repair (BER) pathway is responsible for replacing these damaged bases (4, 5). Within the BER, the damaged base(s) and the corresponding part of the backbone are removed creating a gap of one or multiple nucleotides in the DNA (6). DNA polymerase beta (Pol β) then binds to the gap and subsequently fills the gap by adding cognate nucleotides to the 3’ end of the primer strand (7, 8).

Pol β is one of the smallest eukaryotic polymerases and belongs to the X-family of DNA polymerases (9). It consists of two domains: a polymerase domain and the aforementioned lyase domain (10). Pol β was shown to have an elongated structure in solution (11, 12). Upon binding to gapped DNA, the lyase domain interacts with the 5’ phosphate on the downstream strand, while the polymerase domain adopts a structure that has been compared to a hand (10). Crystal structures have suggested that Pol β bends its DNA substrate with an angle of ~90° (13). Incoming nucleotides then bind to a subdomain known as the “fingers”, forming the ternary complex. A conformational change called “fingers closing” positions the nucleotide closer to the active site to facilitate chemistry. Studies on *E. coli* DNA polymerase I Klenow Fragment (KF), which undergoes a very similar conformational change from an open to a closed conformation, suggested that the fingers do not close entirely when a non-complementary nucleotide is bound. Instead, an intermediate “ajar” conformation was identified, which serves as a “fidelity checkpoint” (14, 15).

In cells, non-complementary nucleotides and ribonucleotides vastly outnumber correct nucleotides; in cancerous cells, nucleotide concentrations can be an order of magnitude higher than in healthy cells (16) further highlighting that effective mechanisms for discriminating correct from non-complementary nucleotides are pivotal for faithful DNA repair. However, for Pol β the existence of additional fidelity checkpoints that can be associated to a ajar conformational state is not widely accepted, although a crystal structure with a mismatched nucleotide suggested that an intermediate conformation might exist (17).

To study fingers movement of Pol β in more detail, Towle-Weicksel *et al.* introduced an assay based on Förster resonance energy transfer (FRET) to monitor fingers-closing using stopped-flow (18). This approach used a fluorescent label on the DNA substrate and a label on Pol β, the latter being attached to the fingers domain after site-directed mutagenesis of valine-303 to cysteine. By fitting their stopped-flow traces to a multi-step kinetic model, the authors were able to extract rates for fingers closing and opening in presence of the complementary nucleotide. Non-complementary nucleotides were not found to induce fingers-closing, leading the authors to hypothesize that discrimination between correct and incorrect nucleotides already takes place before fingers-closing. In later work, the authors showed that a low-fidelity Pol β mutant found in cancer cells exhibits altered fingers dynamics (19).

Here, we developed two single-molecule assays to study the DNA binding behaviour and fingers movement of Pol β, for which we used a combination of FRET and total internal reflection fluorescence (TIRF) microscopy allowing the monitoring of hundreds of molecules in parallel and in real-time. The first assay uses a doubly-labelled gapped DNA substrate to report on binding of unlabelled, wild-type Pol β (wt Pol β). A second assay, inspired by single-molecule work on *E. coli* DNA Pol I by Evans *et al.* (20), employs a similar design as the stopped-flow experiments discussed above: the fingers domain of Pol β is labelled with an acceptor dye, whereas a gapped DNA substrate bears the donor fluorophore. The labelling position on the DNA was chosen such that open and closed conformations of the fingers exhibit different FRET efficiencies (E) when Pol β is bound to the surface-immobilized DNA substrate. This approach allowed us to visualize fingers movement of individual DNA polymerases, which we used to study the response of the polymerase to both complementary and non-complementary nucleotides added to the buffer. We found that non-complementary nucleotides and complementary ribonucleotides lead to partial fingers closing, while complementary nucleotides induce complete fingers closing. This finding supports the idea that a partially closed “fidelity checkpoint” is a more widely employed mechanism by DNA polymerases than initially anticipated as it can be found in both bacterial and eukaryotic DNA polymerases.

## Material And Methods

### Polymerase purification and labelling

In the following, we use the term “wild-type Pol β” to refer to Pol β bearing the substitutions C239S, C267S and V303C, introduced to have a single cysteine residue on the fingers subdomain that can react with the fluorophore bearing a maleimide moiety (18). For the assays in which the fingers conformational change is studied, the V303C was labelled with Alexa Fluor 647 following procedures described before (18). The labelling efficiency was 60–70% as determined by SDS-polyacrylamide gel electrophoresis (data not shown). For experiments with *E. coli* DNA Polymerase I Klenow Fragment, we used the D424A mutant (hereafter referred to as simply “KF”) that abolishes the 3’ to 5’ exonuclease activity of KF.

### DNA substrate design

As a first step to construct a gapped DNA construct labelled at adequate positions with fluorophores, we examined crystal structures 1BPX and 1BPY (13) that represent Pol β bound to gapped DNA with open and closed fingers, respectively. We extended the DNA from the crystal structures on both sides of the polymerase with a B-DNA helix of our preferred sequence using the 3D-DART server (21). Next, we used FPS (short for *FRET-restrained positioning and screening)* software to model the accessible volumes of the fluorophores at potential labelling positions on the DNA and determine inter-dye distances <RDA>_*E*_ (22). Modelling parameters include the dimensions of the fluorophore and the dimensions of the linker (Supplementary Table 1). We selected two labelling positions (−15 and +12, see Fig. 1A-C) that are located outside the binding region of Pol β. Using Cy3B as a donor fluorophore on the −15 position and Cy5 as an acceptor on +12, these positions are within the distance range for FRET (*R_o,cy3B→cy5_* = 6.9 nm, <RDA>*_E,model_* = 6.0 nm). Additionally, the Cy3B on the - 15 position is also close enough to the fingers subdomain to exhibit FRET with the Alexa Fluor 647 (*R_0,Cy3B→Alexa Fluor 647_* = 6.9 nm, <RDA>*_E,fingers open_* = 6.4 nm, <RDA>*_E,ftngers closed_* = 5.5 nm, Fig. 1D-E). Importantly, these distances translate to a large difference in FRET efficiency (*E*) between the open (*E* = 0.60) and closed (*E* = 0.79) conformations of the fingers.

**Figure 1.**
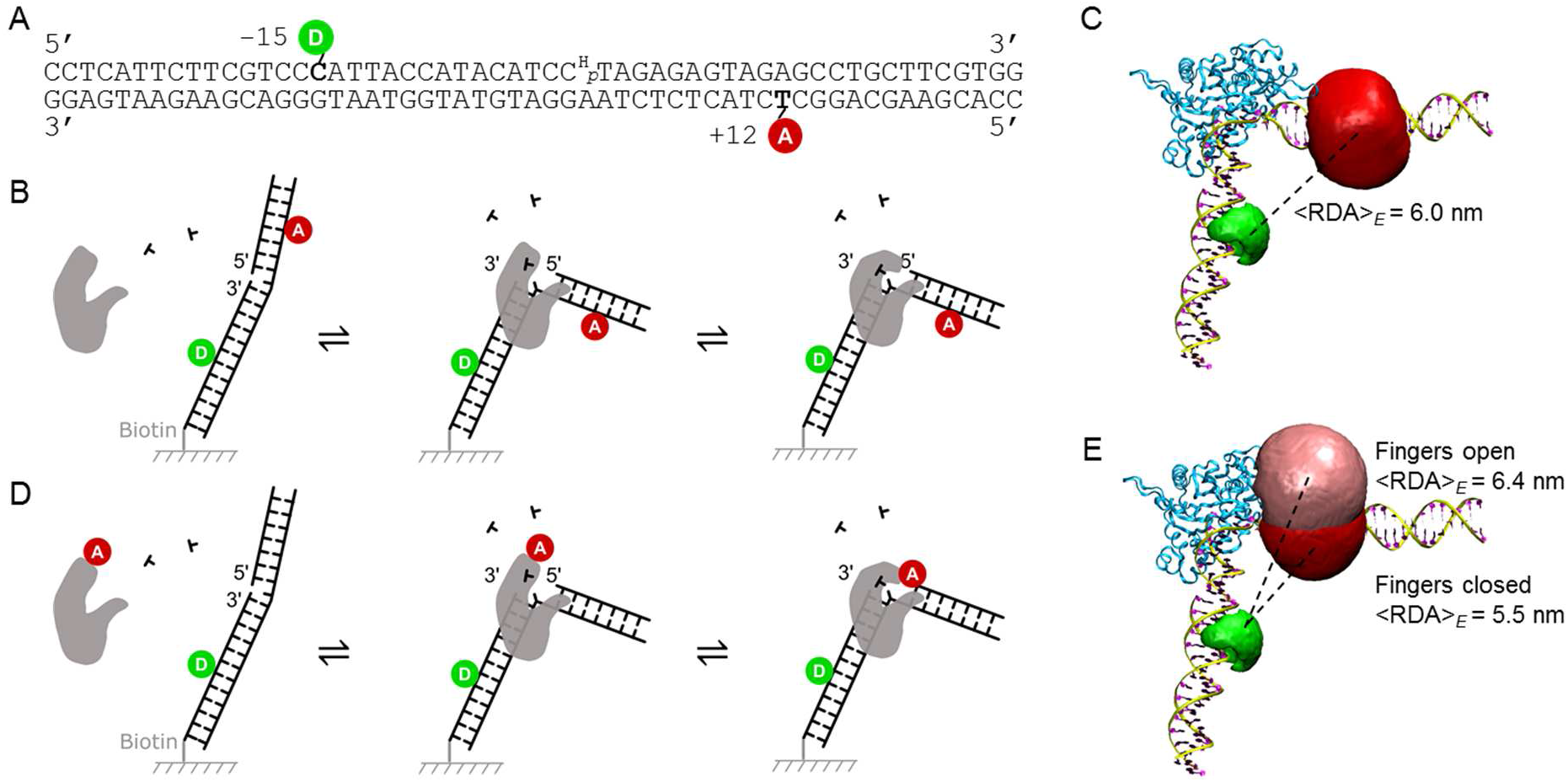
Experimental design. **A)** Sequence and labelling positions of the gapped DNA construct. The 3’ end of the primer is dideoxyterminated and a phosphate group is present at the 5’ end of the complementary strand. **B)** Schematic overview of the assay with the bending sensor. A gapped DNA substrate is thought to bend upon binding of Pol β, which translates to a change in FRET efficiency. **C)** Extended crystal structure (pdb: 1BPX). Accessible volumes of Cy3B (green) and Cy5 (red) are outside the putative binding region of the polymerase. **D)** Schematic overview of the “fingers closing” assay. Pol β, labelled with Alexa Fluor 647 on the fingers domain, binds to a gapped DNA substrate that is labelled with Cy3B. Fingers movement results in a change in FRET efficiency. **E)** Structure 1BPX with the accessible volumes of Alexa Fluor 647, in the open (pink) and closed (red) conformation. (The accessible volume of the closed conformation was modelled using structure 1BPY, which is not shown here.) Cy3B is on the primer. The distance between the donor and acceptor dyes decreases with fingers closing.

We annealed the 1 nt gapped DNA complex using a template A from a 30-mer primer sequence (biotin-5’-CCT CAT TCT TCG TCC CAT TAC CAT ACA TCCH-3’), a 55-mer template sequence (5’-CCA CGA AGC AGG CTC TAC TCT CTA AGG ATG TAT GGT AAT GGG ACG AAG AAT GAG G-3’) and a 24-mer downstream complementary strand (5’-phosphate-TAG AGA GTA GAG CCT GCT TCG TGG-3’), which we ordered from IBA Life Sciences (Germany) and Eurogentec (Belgium). All oligos were HPLC or gel purified prior to use. The primer sequence was internally labelled with donor dye Cy3B through a C6 linker at the previously determined −15 cytosine base; for experiments with unlabelled wt Pol β, also the template was internally labelled with Cy5 through a C6 linker at the +12 thymine base.

### TIRF experiments

Labelled DNA molecules were immobilized on PEGylated glass coverslips using a protocol described before (20). We used flow channels formed by Ibidi sticky-Slides VI^0.4^. Molecules were imaged on a home-built TIRF microscope, described in more detail elsewhere (23). All experiments were performed using Alternating Laser Excitation (ALEX) in which the direct excitation of the donor alternates with the direct excitation of the acceptor fluorophore (24, 25). Experiments on wt Pol β and doubly labelled DNA were performed with laser powers of 1.5 mW (λ = 561 nm) and 1.5 mW (λ = 638 nm). Excitation time and camera frame time were set to 50 ms. Raw FRET efficiency (*E\**) was calculated using *E\** = *DA* / (*DD* + *DA*), in which *DD* is donor emission intensity after donor excitation, and *DA* is acceptor emission intensity after donor excitation (FRET). Acceptor emission intensity after acceptor excitation *AA*, as obtained during ALEX, was used for time trace selection. Experiments with fluorescently labelled POL B were performed with laser powers of 1.5 mW (λ = 561 nm) and 0.75 mW (λ = 638 nm). Excitation time and frame time were 25 ms. Surface-immobilized DNA molecules were imaged in a buffer containing either wt Pol β (10, 30, 100 and 300 nM) or labelled Pol β-V303C-Alexa Fluor 647 (10 nM). Imaging buffer further contained 50 mM Tris (pH 7.5), 10 mM MgCl_2_, 100 mM NaCl, 100 μg/mL BSA, 5% glycerol, 1 mM DTT, 1 mM Trolox, 1% gloxy and 1% glucose. Trolox is a triplet state quencher (26); gloxy and glucose form an enzymatic oxygen scavenger system to prevent premature fluorophore bleaching (27). When specified, complementary dTTPs were added to achieve final concentrations of (0.1, 0.5, 1, 2, 5, 10 and 50) μM; concentrations of non-complementary dGTPs and rUTPs were (10, 30, 100, 300, 1000 and 3000) μM. For the experiments with KF, an imaging buffer without NaCl was used.

### Time trace selection and Hidden Markov Modelling

Time traces from individual molecules were collected to measure polymerase binding times and to extract dwell times of open and closed fingers conformations. Because of variations in the signal-to-noise ratio of molecules as well as the presence of bleaching and blinking, an initial selection of molecules was made by hand: only DNA molecules that showed a constant *DD*+*DA* signal with sudden transitions (within 1 frame) from the free to the bound state were selected. To determine the binding times, we first applied a 5 frame moving median filter to all selected traces, before applying additional selection criteria: 1) the sum of *DD* and *AA* is higher than 50 photons and 2) the FRET efficiency is higher than 0.4. Additionally, settings were such that disappearance of donor signal (bleaching) was interpreted as the end of the trace, and disappearance of acceptor signal (bleaching or polymerase dissociation) was interpreted as the end of a binding event. Filtering traces following these criteria sometimes resulted in longer binding events being cut in multiple shorter events due to remaining noise. To prevent these cases, an exception was added to allow single point excursions to lower intensities or FRET efficiencies. The final algorithm was found to identify most binding events that are also detectable by visual inspection. Extremely short events, however, are often not detected because of the strong median filter. These events may therefore be under-represented in our dwell time histograms.

For extraction of fingers conformational changes, binding events from experiments with dTTPs were loaded into ebFRET, a software package for Hidden Markov Modelling (28). Because the final point of some binding events may have a donor or acceptor that is already decreasing in intensity (just before the cut-off that we set for dissociation or bleaching), we removed these points from the traces by applying a padding of 10 time points. Next, the prior for the minimal centre position (open fingers) was set to *E\** = 0.4, and that for the maximal centre position (closed fingers) to *E\** = 0.8. The convergence threshold was set to 10^−5^.

## Results

### Pol β bends its substrate, but in a different fashion than crystal structures suggest

First, we assessed binding of wt Pol β to dsDNA with a 1 nt gap, mimicking the BER pathway intermediate that is the natural and preferred substrate of Pol β (10). Crystal structures 1BPX and 1BPY suggested that the DNA adopts a sharply bent conformation (~90°) after binding of Pol β. Bending of the DNA is therefore an indicator for polymerase binding. We labelled our DNA substrate with a donor dye on the primer and an acceptor dye on the template, at positions that are outside the binding region of the polymerase (as judged from crystal structures 1BPX and 1BPY), thus creating a “bending sensor”.

The native DNA substrate has a FRET efficiency *E\** = 0.37 (Fig. 2A) which corresponds to an inter-fluorophore distance of 7.8 nm after correcting for leakage, direct excitation and gamma (25) (see Supplementary Table 2 for step-by-step correction). A model of a rigid double-stranded DNA helix in PyMOL predicted an inter-dye distance <RDA>_E_ of 8.3 nm. These values are slightly different, but we note that our static DNA model cannot account for the flexibility introduced by the gap.

**Figure 2.**
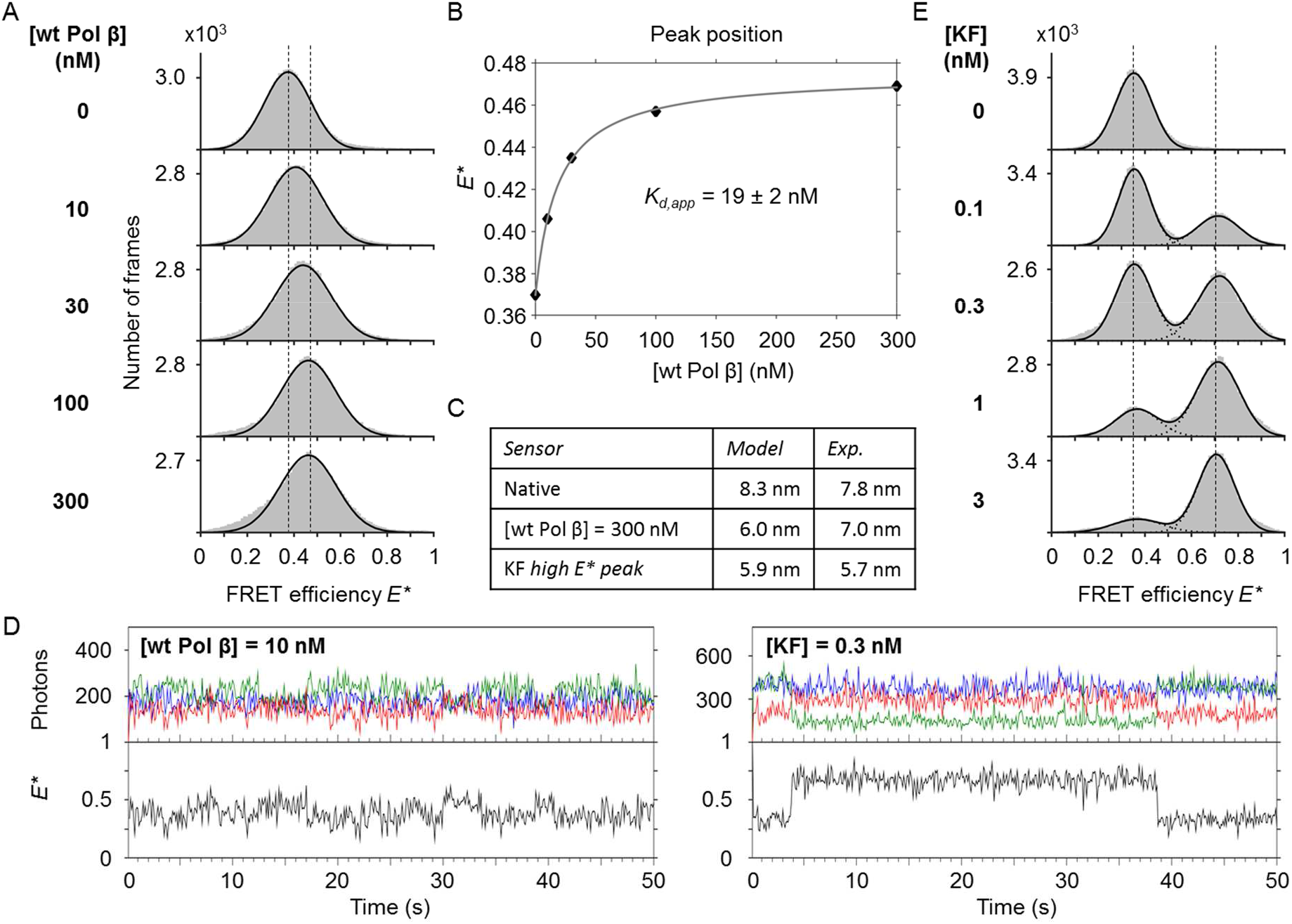
Response of the gapped DNA bending sensor to wt Pol β and KF. **A)** The mean FRET efficiency *E\** increases with increasing concentrations of wt Pol β. **B)** Peak positions from A) plotted against wt Pol β concentration, fitted to a Langmuir binding isotherm (grey line) reveal an apparent *Kd* of 19 ± 2 nM. **C)** Modelled and experimentally determined inter-dye distances of both the native bending sensor and the bent conformation. **D)** Time traces of a single DNA bending sensors at [wt Pol β] = 10 nM and [KF] = 0.3 nM. **E)** Bending sensors respond differently to increasing concentrations of KF, showing a growing occupancy of a new high-FRET species.

Increasing the concentration of Pol β leads to a peak shift, levelling off towards a FRET efficiency of ~0.47 (Fig. 2B), reflecting the bent conformation of the DNA and, after corrections, a distance of 7.0 nm (Fig. 2C). The distance calculated from structures 1BPX and 1BPY is significantly shorter (6.0 nm). We used the peak shift to determine the dissociation constant *K_d_*, describing the binding affinity of DNA and Pol β, and obtained *K_d_* = 19 ± 2 nM, in good agreement with previously published values (19, 29). A trace of a single DNA molecule at [wt Pol β] = 10 nM shows excursions between the low and high FRET states (Fig. 2D). The interpretation, however, is complicated by the noise in combination with a small Δ*E*\*.

Additionally, we tested the response of our bending sensor to *E. coli* DNA Polymerase I Klenow Fragment (KF). Like Pol β, this polymerase bends a gapped DNA substrate to an angle of approximately 90° (as determined by FRET measurements and structural modelling by Craggs *et al.* (BioRxiv: https://doi.org/10.1101/263038)). Upon increasing the concentration of KF, we did not observe a peak shift; instead, we saw a new peak appearing at *E\** = 0.71 (Fig. 2E). Accurate FRET calculations revealed a corresponding inter-fluorophore distance of 5.7 nm (Fig. 2C, Supplementary Table 2) in excellent agreement with a FRET restrained structural model by Craggs *et al.* predicting a distance of 5.9 nm.

### Hidden Markov Modelling resolves fingers movement in presence of complementary dTTPs

Next, we studied the ability of labelled Pol β to report on the conformation of the fingers subdomain. We performed a titration of labelled Pol β with increasing [A-dTTP] concentrations. Time traces of single DNA molecules showed binding events of labelled Pol β as an increase in *AA* and the appearance of FRET (Fig. 3A). Hidden Markov Modelling (HMM) was used to identify the open (low *E\**) and closed (high *E\**) conformations within time traces of individual binding events (Fig. 3B). At [dTTP] = 0.5 μM, traces predominantly show low FRET efficiency, with only short excursions to the high FRET efficiency that is associated with closed fingers. At higher dTTP concentrations, longer residence times in the closed state are observed. We constructed FRET efficiency histograms and indicated the open and closed populations, as determined by HMM (Fig. 3C). Without dTTPs being present, the fingers mostly adopt the open conformation (92%). With increasing dTTP concentration, the closed conformation is increasingly populated. At [dTTP] = 50 μM, the fingers are mostly closed (95%).

**Figure 3.**
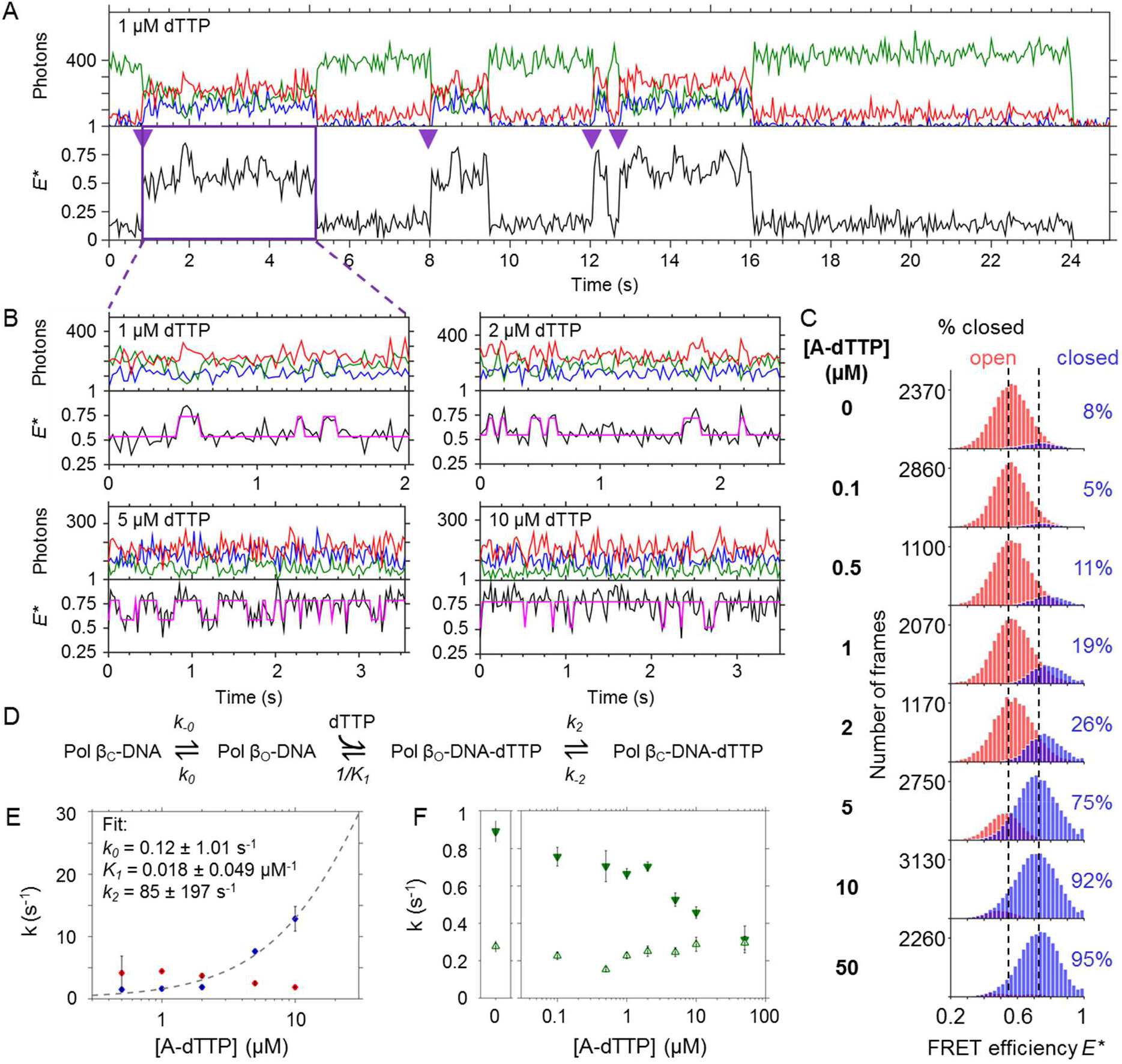
Fingers opening and closing of Pol β revealed by smFRET. **A)** Time trace of a single DNA molecule in presence of 10 nM labelled Pol β and 1 μM complementary dTTPs. Pol β binding events are indicated with purple triangles. At *t* = 24 s, the donor bleaches. **B)** Time traces of labelled Pol β - DNA complexes, at various concentrations of dTTP. The first trace ([dTTP] = 1 μM) is a binding event taken from A). FRET efficiency *E\** (black trace) is calculated from the *DD* signal (green trace) and the *DA* signal (red trace) in the upper panel. The *AA* signal (blue trace) is shown here as well to indicate that the observed events are not due to acceptor photophysics. An HMM fit (magenta) indicates the open and closed conformation of the fingers. **C)** Corresponding FRET efficiency histograms (32 bins between *E\** = 0.2 and 1) of the Pol β - DNA complexes. The FRET efficiencies of the open (shown in red) and closed (shown in blue) conformation were plotted after the states have been assigned via HMM. **D)** Schematic model used to describe the dynamics of fingers movement in the binary (k_o_/k_-o_) and ternary complex (*k_2_/k_-2_*). **E)** The observed closing rates *k_close,obs_* (blue) and opening rates *k_open_* (red) plotted against [dTTP]. Data were fit to a function described in the main text (dashed line) and derived from the model depicted in C). Error bars represent the 95% confidence interval obtained from fitting the dwell times. **F)** Complementary dTTPs stabilize the Pol β - DNA complex. Rates *k_on_* (green open triangles) and *k_off_* (green solid triangles) plotted against [dTTP]. *k_off_* decreases with increasing concentrations of dTTP, while *k_on_* remains constant. Error bars represent the 95% confidence interval obtained from fitting the dwell times.

Using accurate FRET, we determined the distances associated with open and closed fingers (Supplementary Table 3). We found an inter-fluorophore distances of 6.5 nm for the open and 5.6 nm for the closed conformation. We note that these distances are in excellent agreement with the distances of 6.4 nm and 5.5 nm as predicted from structural modelling with FPS software (see Methods).

Dwell time histograms of the open and closed conformations were constructed and fitted with exponential decay curves (concentrations of 0, 0.1 and 50 μM were left out due to an insufficient number of transitions). The rate of fingers closing is extracted from the dwell times in the open conformation and vice versa. Plotting *k_obs,close_* and *k_open_* against the concentration of dTTPs showed that the closing rate is concentration-dependent; the opening rate varies as well, but to a lesser extent. A simple model that links fingers closing to the affinity of complementary nucleotides was previously described (Fig. 3D) (20). Here, the concentration of dTTPs relates to *k_obs,close_* as follows:

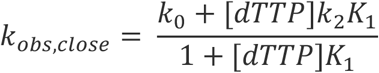

in which *k_0_* is the closing rate of the binary complex, *K_1_* is the association constant for dTTPs and *k_2_* is the closing rate with dTTP bound to the fingers. As fingers closing was very rare in our traces at [dTTP] = 0 μM, we could not provide the model with a value for *k_0_*. We therefore fitted our data with no constraints for *k_0_*, *K_1_* and *k_2_* (Fig. 3E) and found a *k_2_* of 85 ± 197 s^-1^ (mean ± standard deviation), *K_1_* of 0. 018 ± 0.049 μM^−1^ (corresponding to a *K_d(dTTP)_* of 56 μM) and a *k_0_* of 0.12 ± 1.01 s^−1^.

We noted that the duration of Pol β - DNA binding events increased with increasing [dTTP] by observing an increase of *k_off_*, while *k_on_* is not affected (Fig. 3F).

### Fingers close only partially in presence of non-complementary dNTPs and complementary rNTPs

To investigate the potential existence of a “partially closed” or “ajar” conformation in Pol β, we studied the fingers conformational change in the presence of non-complementary dNTPs and complementary rNTPs. Previous work has shown that increasing concentrations of an incorrect nucleotide shift the position of the “fingers open” peak of KF towards a slightly higher FRET efficiency, likely caused by the polymerase quickly screening and rejecting incorrect nucleotides (15, 20). For Pol β, we performed a similar experiment: we tested a range of different concentrations of dGTPs and rUTPs, and plotted the position of the main peak (Fig. 4A-C). Indeed, for both incorrect nucleotides a shift in *E\** from ~0.55 to ~0.59 is observed, suggesting that Pol β (like KF) has a fidelity checkpoint at a partially closed fingers conformation. The associated Δ*E\**, however, is too small for clear detection of this state in individual single-molecule time traces.

**Figure 4.**
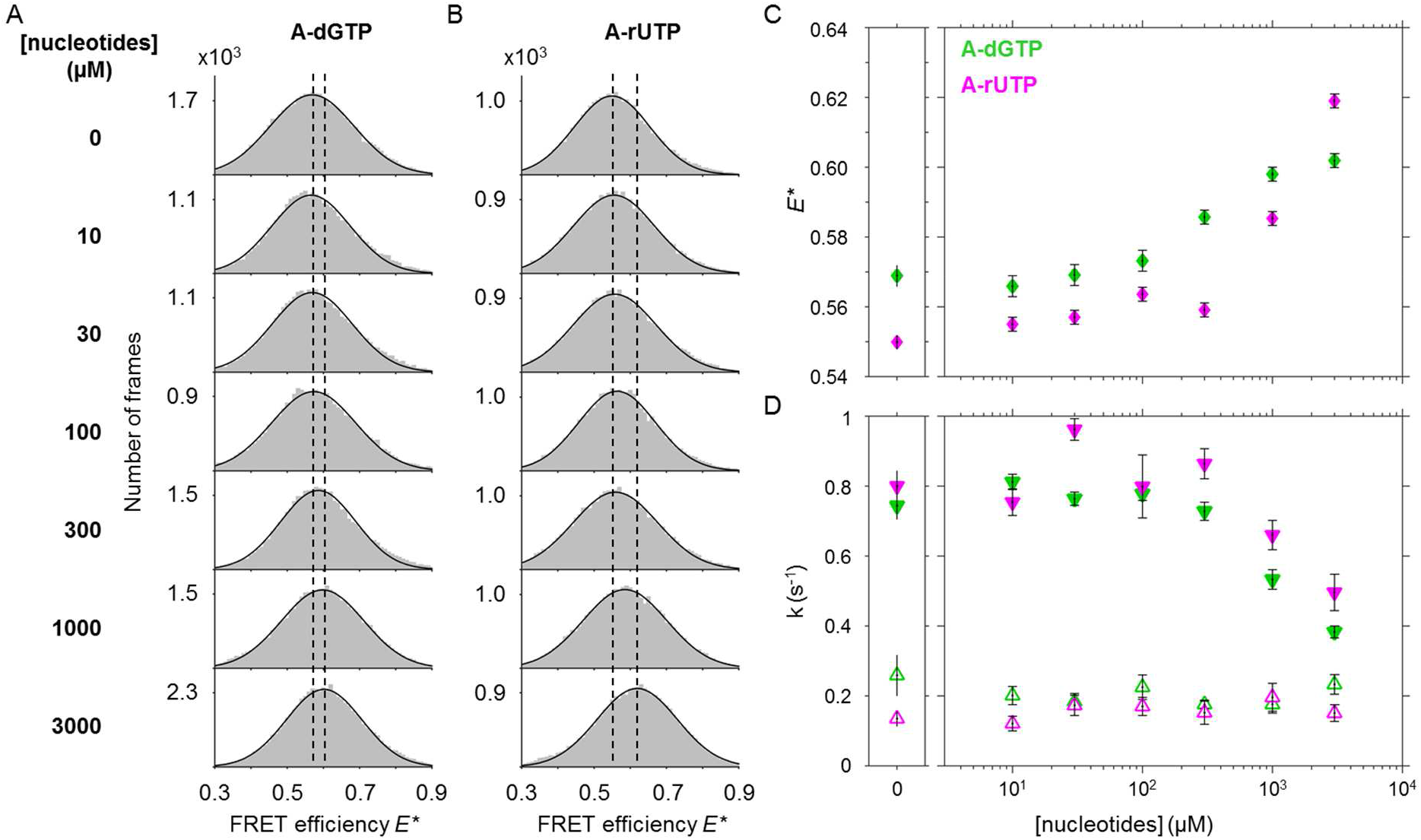
Incorrect nucleotides lead to partial fingers closing and stabilize the ternary complex. **A)** FRET efficiency histograms of Pol β-DNA complexes at increasing [A-dGTP]. Histograms were filtered based on colocalization of donor (DD) and acceptor (AA) fluorophores (see Supplementary Information). Dashed lines are added for visual guidance. **B)** FRET efficiency histograms of Pol β - DNA complexes at increasing [A-rUTP]. **C)** Position of the main peak plotted against [nucleotide]. Both dGTPs (green diamonds) and rUTPs (magenta diamonds) cause a shift in the peak of the open conformation. Error bars indicate the 95% confidence intervals on the fits. **D)** Rates *k_on_* (open triangles) and *k_off_* (solid triangles) at increasing concentrations of dGTPs (green) and rUTPs (magenta). Error bars indicate the 95% confidence intervals on the fits. *k_off_* decreases with increasing concentrations of incorrect nucleotides, while *k_on_* remains constant.

We asked if incorrect nucleotides have an influence on the stability of the ternary complex. Evans *et al.* and Markiewicz *et al.* showed that non-complementary nucleotides increase *k_off_* for KF (20, 30). We identified all polymerase binding events in our time traces with labelled Pol β and constructed dwell time histograms. Fitting with an exponential decay function yielded values for *k_on_* and *k_off_*for every nucleotide concentration of the titration series (Fig. 4D). Interestingly, *k_on_* and *k_off_* remained constant over a wide range of the nucleotide concentration series, corresponding with earlier findings that the interaction of the lyase domain with the 5’-phosphate plays a large role in substrate binding (31). At high nucleotide concentrations, however, *k_off_* decreased, indicating that for Pol β bound to gapped DNA even incorrect nucleotides stabilize the ternary complex albeit only at higher concentrations than for the correct dNTP.

## Discussion

The use of smFRET allowed us to observe and analyse conformational changes of individual Pol β - DNA complexes in real time, thereby overcoming some of the ensemble averaging inherent to conventional fluorescence-based techniques, such as stopped-flow.

Our experiments with wt Pol β showed substrate binding with a *K_d_* of 19 ± 2 nM. Other studies have also reported values in the low nanomolar range for gapped DNA constructs: fitting stopped-flow data resulted in a *K_d_* of 5 nM (19), and a titration based on single-turnover analysis at different DNA concentrations revealed a *K_d_* of 22 nM (29). Surprisingly, strong DNA bending upon polymerase binding, as predicted by various structures resolved with X-ray crystallography, does not occur to the expected extent suggesting that the bend angle in solution is larger than 90°. Compared to conditions required for successful crystallisation, we were able to use a DNA sequence with longer up - and downstream strands (here: upstream region 30 base pairs, downstream region 24 base pairs; pdb: 1BPX is 10 and 5 base pairs, respectively). While an upstream binding region of 11 ± 1 nt was revealed with fluorescently labelled DNA previously (32), the short downstream region in 1BPX raises the question whether the crystal structures obscure interactions of the polymerase with parts of the DNA that are farther away from the gap.

We further studied the conformational change associated to “fingers closing” using a fluorescently labelled version of Pol β. We observed an increase in the rate of fingers closing with increasing concentrations of the complementary dNTP, as expected for an induced fit mechanism. Previous studies (18, 19) found a rapid fingers closing rate of 98 s^−1^, close to the maximum rate of 85 s^−1^ that we determined from our fit. The opening rate is concentration-dependent as well, and decreases with increasing concentrations of dTTPs. This may indicate that *k_-2_* is lower than *k_-o_*, i.e. the closed state is stabilized when a complementary nucleotide is bound. However, we note that the likelihood of missing events increases with dTTP concentration, leading to lengthening of the closed-state dwell times and extraction of a lower apparent opening rate. Generally, because of our time resolution of 50 ms, the dynamics of opening and closing become increasingly difficult to analyse at dTTP concentrations higher than 10 μM.

The *K_d_*of the incoming nucleotide (1 / *K_1_* = 56 μM) is higher than determined before using stopped flow and chemical quench analysis (*K_d_* = 2.5 μM) (18, 19). The reason for this difference is that we fitted our observations to a model that accounts for fingers opening and closing in presence and absence of dNTPs. An analysis based on the assumption that fingers close only and immediately after nucleotide binding, however, would find a *K_d_* at the concentration where the open and closed conformations are equally populated (in our data: between 2 and 5 μM).

We found evidence that the fingers adopt a partially closed conformation when supplied with incorrect nucleotides, reminiscent of KF. Both addition of dGTPs and rUTPs result in partial closing, indicating that at this ‘fidelity checkpoint’, incoming nucleotides are both screened for complementarity and backbone structure. Ensemble studies on KF suggested that complementary rNTPs proceed farther in the reaction pathway than non-complementary dNTPs (32, 33), but the difference in *E\** is too small in our assay to investigate that further. A crystal structure of Pol β with a dG-dAMPCPP mismatch in the active site initially suggested that fingers close at least partially (17). It will be interesting to see in follow-up single-molecule studies whether this checkpoint is more dominantly populated in mutator variants of Pol β, as it has been shown for bacterial DNA Pol I (15).

Furthermore, we found that non-matching nucleotides do not destabilize the polymerase-DNA complex, as was found to be the case for KF (20, 30). Instead, at high concentrations (> 1 mM) these nucleotides stabilize the Pol β - DNA complex by lowering *k_off_*. This seems counterintuitive since prolonged binding times of a flawed ternary complex may increase the chance of actual incorporation of the wrong base. Non-complementary nucleotides, however, are always more abundant in cells than the correct dNTPs. Considering nucleotides are randomly sampled, repair of damaged DNA may take much longer if a ternary complex with a non-complementary nucleotide is destabilized.

The main advantage of using smFRET to study fingers movement is the direct observation of this conformational change in individual molecules, allowing to calculate rates for opening and closing. In addition, smFRET provides access to the associated intermolecular distances, which support our interpretation that the observed fluctuations in FRET efficiency are indeed switching between an open and closed state. An even more direct method to study fingers opening and closing would see both fluorophores on the polymerase, to avoid convolution of fingers movement with any potential movement in the DNA substrate.

## Supplementary Data

Supplementary Data is available online.

## Acknowledgement

*E. coli* DNA Polymerase I Klenow Fragment was kindly supplied by Catherine Joyce, Tim Craggs and Achillefs Kapanidis.

